# A smartphone-based tool for rapid, portable, and automated wide-field retinal imaging

**DOI:** 10.1101/364265

**Authors:** Tyson N. Kim, Frank Myers, Clay Reber, PJ Loury, Panagiota Loumou, Doug Webster, Chris Echanique, Patrick Li, Jose R. Davila, Robi N. Maamari, Neil A. Switz, Jeremy Keenan, Maria A. Woodward, Yannis M. Paulus, Todd Margolis, Daniel A. Fletcher

**Affiliations:** Department of Ophthalmology and Visual Sciences, University of Michigan School of Medicine; Department of Bioengineering and Biophysics Program, University of California, Berkeley; Department of Ophthalmology, University of California, San Francisco; Department of Ophthalmology and Visual Sciences, Washington University School of Medicine in St. Louis; Department of Physics and Astronomy, San José State University; Chan-Zuckerberg Biohub, San Francisco, CA

**Author notes:** Authors have contributed equally to this work (Tyson N. Kim, Frank Myers, Clay Reber). **Grant information:** This work was supported in part by the Michigan Translational Research and Commercialization program with support from MEDC and the UM Medical School (TNK), NEI grant no. K23 EY023596 (MAW), QB3 Bridging the Gap Awards from the Rogers Family Foundation (DAF) and a Bakar Fellows Award (DAF). DAF is a Chan-Zuckerberg Biohub investigator.

## Abstract

**Purpose:** High-quality, wide-field retinal imaging is a valuable method to screen preventable, vision-threatening diseases of the retina. Smartphone-based retinal cameras hold promise for increasing access to retinal imaging, but variable image quality and restricted field of view can limit their utility. We developed and clinically tested a smartphone-based system that addresses these challenges with automation-assisted imaging.

**Methods:** The system was designed to improve smartphone retinal imaging by combining automated fixation guidance, photomontage, and multi-colored illumination with optimized optics, user-tested ergonomics, and touch-screen interface. System performance was evaluated from images of ophthalmic patients taken by non-ophthalmic personnel. Two masked ophthalmologists evaluated images for abnormalities and disease severity.

**Results:** The system automatically generated 100-degree retinal photomontages from five overlapping images in under 1 minute at full resolution (52.3 pixels per retinal degree) fully on-phone, revealing numerous retinal abnormalities. Feasibility of the system for DR screening using the retinal photomontages was performed in 71 diabetics by masked graders. DR grade matched perfectly with dilated clinical examination in 55.1% of eyes and within 1 severity level for 85.2% of eyes. For referral-warranted DR, average sensitivity was 93.3% and specificity 56.8%.

**Conclusions:** Automation-assisted imaging produced high-quality, wide-field retinal images that demonstrate the potential of smartphone-based retinal cameras to be used for retinal disease screening.

**Translational Relevance:** Enhancement of smartphone-based retinal imaging through automation and software intelligence holds great promise for increasing the accessibility of retinal screening.

## Introduction

Retinal photography is used extensively to assist with diagnosis and monitoring of retinal diseases. However, access to traditional tabletop retinal cameras is limited by their high cost, bulky design, and need for skilled operators. Conventional imaging approaches also require patient cooperation with stabilized, upright head positioning, which can be difficult among sick, wheelchair-bound, and immobilized patients, as well as children^1^. Handheld ophthalmoscopes with digital image capture offer an alternative. Recently, smartphone-based retinal imaging approaches have been described that leverage the phone for its portability, high-resolution camera, and wireless data transfer capabilities to capture diagnostic images for real-time or remote anterior^2^ or posterior segment^3–12^ evaluation at a fraction of the cost of traditional instruments^8–10, 12^. These investigations nicely demonstrate the growing potential for smartphone imaging to expand the accessibility of ophthalmic care and photographic screening of vision threatening diseases.

There is particularly great interest in the validation and integration of smartphone-based retinal photography in local community screening programs for diseases such as glaucoma and diabetic retinopathy (DR)^13–16^, as well as in the emergency department and inpatient settings where the fundus examination is under-performed^17–19^. DR is the most common microvascular complication of diabetes and the leading cause of vision loss in working-age adults aged 20 to 74 in the United States, accounting for 12% of new cases annually^20–23^. The World Health Organization estimates more than 346 million people worldwide have diabetes and that 552 million people will be affected by 2030^24^. Physicians have effective treatments for complications of DR, and it has been shown that patients have better vision outcomes with early detection and treatment^25–27^. However, despite well-accepted guidelines^28^, nearly half of diabetic adults in the US do not receive recommended annual screening for DR, and vulnerable populations with less access to specialty medical care have an estimated screening rate between 10-20% per year^29^. Ophthalmologists recognize that telehealth screening of DR via retinal photography in the primary care setting, with remote ophthalmologist consultation^30^, is a mechanism to improve access thereby improving outcomes in a cost-effective manner^31, 32^. A remaining challenge is how to increase access to such retinal photography.

The gold-standard photographic screening technique for DR is 7-field, 30-degree mydriatic tabletop retinal photography^33^ in which 14 images per eye from 7 standard fields comprising the posterior 90 degrees of the retina are evaluated to determine the risk of vision loss and retinopathy progression. Subsequent studies have demonstrated that 3-field, 45-degree non-mydriatic fundus photography was effective in grading DR to determine referral-warranted disease, whereas a single 45-degree photograph was insufficient to accurately grade DR^34–36^. Importantly, imaging by non-expert operators may affect image quality and is an important consideration for community-based screening efforts^37, 38^. Several studies have investigated smartphone-based screening of DR^4, 16, 39^ and reported sensitivities of ≥80% with high specificities, as recommended by the British Diabetic Association for new imaging devices in population-based screening^40^. However, other investigators have found insufficient sensitivities of below 60% for detecting DR^37^ suggesting smartphone imaging is not universally reliable. In light of this, smartphone-based retinal imaging has continued to undergo validation against more traditional photographic methods^37, 41–43^.

We previously reported on the development of the Ocular CellScope, a handheld, smartphone-integrated imaging device capable of capturing high-quality, wide-field images of the retina[3]. From extensive field-testing with that device, we identified several challenges facing the performance and use of that device in particular and smartphone-based retinal photography in general. First, surveying wide regions of the retina with a handheld device is technically challenging and requires extensive operator experience to comprehensively image the peripheral retina. Second, sustained levels of bright white-light illumination for high-resolution images of the retina can be uncomfortable for patients, resulting in poor image localization caused by gaze shifts and decreased image quality from motion artifacts. Third, smartphone imaging approaches have typically required two-handed operation and been asymmetric, with different orientation for the right and left eye, increasing operator instability and motion. Fourth, the lack of streamlined data management for acquiring, viewing, and storing images slows workflow and increased examination time.

Here, we demonstrate a retinal imaging system, CellScope Retina, that addresses these issues by incorporating automation to improve image quality and reliability. We describe the application of CellScope Retina in feasibility studies of inpatient and outpatient settings, where images of various retinal pathologies, including diabetic retinopathy, were captured. The 100-degree field was chosen to ensure that a larger portion of the retina is imaged with CellScope Retina than the gold-standard Early Treatment Diabetic Retinopathy Study (ETDRS) screening technique, which evaluates the posterior 90 degrees using seven individual 30-degree field images. Additionally, CellScope Retina images have a resolution of 52.3 pixels per retinal degree, surpassing the minimum resolution requirement of 30 pixels per degree described by the National Health Service for diabetic retinopathy^44^. To our knowledge, CellScope Retina is the first demonstration of an automated smartphone-based system capable of imaging, stitching, and reviewing a wide-field retinal montage in a fully handheld platform without requiring an external computer.

### Methods

#### Hardware Design

The CellScope Retina weighs ~310 grams and consists of a smartphone and 3D-printed plastic housing containing optics for illuminating and imaging the retina onto the camera of a smartphone (Fig. 1A). A polarizing wire grid beam splitter (Moxtek HCPBF02A, Orem, UT) is used to illuminate the retina with polarized light and minimize unwanted reflections as previously described[3]. A custom printed circuit board (PCB) provides illumination with a single white LED (Lumileds Luxeon LXML-PWN2, Brantford Ontario, Canada) at the center and a ring of radius 3.15 mm with three 655nm deep red LEDs (Lumileds Luxeon LXM3-PD01, San Jose, CA). An aspheric condenser lens (Thorlabs ACL12708U-A, Newton, NJ) transmits light from the LEDs to a diffuser (Thorlabs ED1-C50), through a polarizer, an annular mask with 8mm inner diameter and 15mm outer diameter, and a 50mm focal length condenser lens (Thorlabs LA1171-A). The illumination light is reflected by a beam splitter and passes through a 54-diopter ophthalmic lens (Ocular Instruments OI54-A, Bellevue, WA) to form an annulus with a 4.8mm inner diameter and 9.6mm outer diameter at the surface of the cornea. The illumination light defocuses and uniformly illuminates the retina, from which it is scattered with accompanying depolarization. Light from the eye is collected and transmitted through a second polarizer (Edmund Optics 85919, Barrington, NJ) thereby preferentially blocking reflected (and orthogonally polarized illumination light. An achromatic lens with a focal length of 20mm (Edmund Optics 47661) relays the signal to the smartphone camera module.

**Figure 1.**
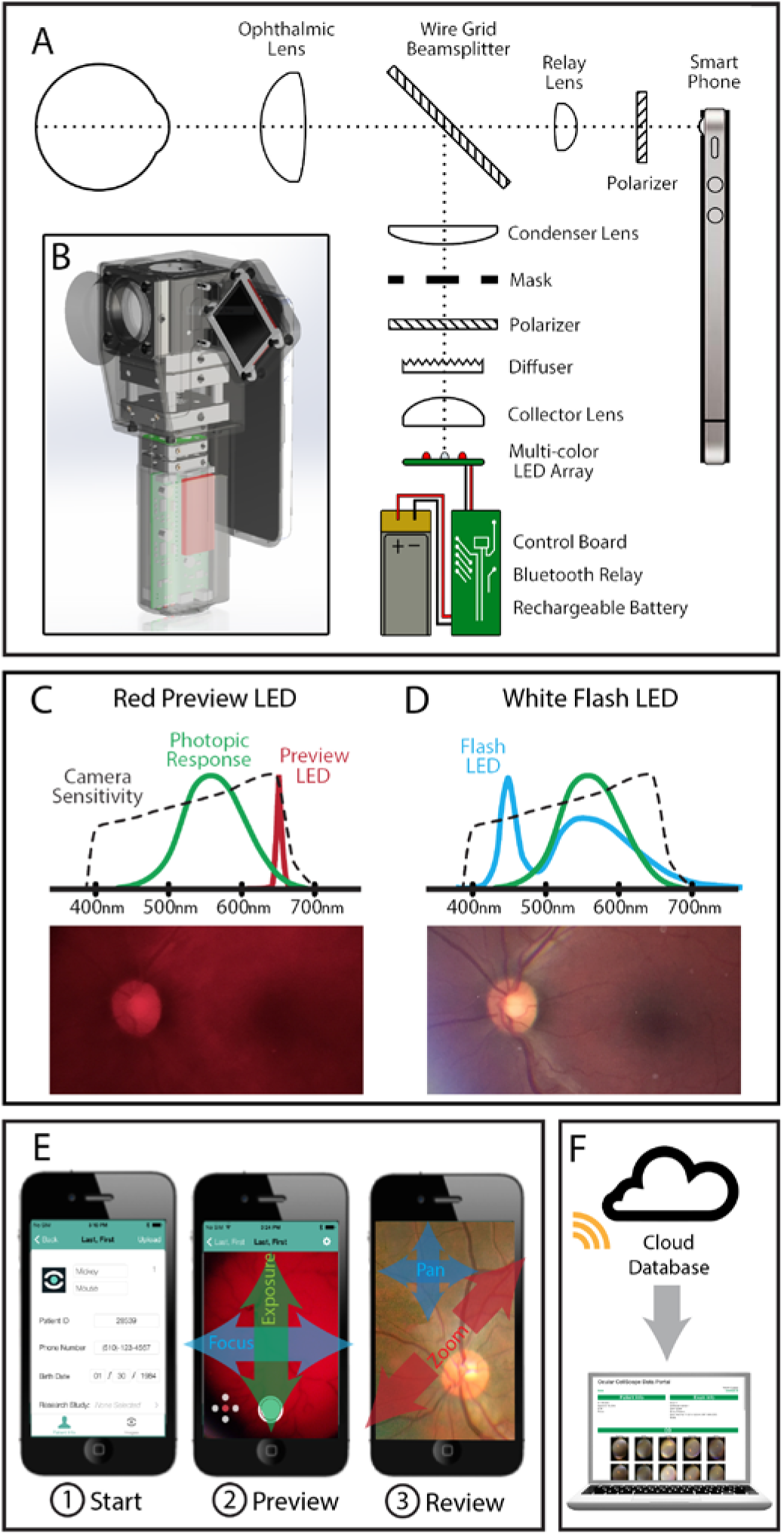
CellScope Retina schematic and workflow. (A) Schematic of the optical system. Light from LEDs is directed through a mask and forms an annulus that passes through the peripheral cornea and focuses through the pupil. In propagating through the eye the light becomes defocused, providing even illumination at the retina. Polarization filters minimize unwanted reflections from anterior ocular surfaces, enabling the smartphone to capture a clear image of the retina. (B) The compact optical system and custom-control electronics fit inside a handheld enclosure. (C) Red LED illumination of 633nm peak emission is used for focusing on the retina which is within the spectral range of the iPhone camera but outside the peak photopic response of the eye. (D) A white LED with a broad emission spectrum is flashed for recording images of the retina. LED spectra in (C) and (D) are from respective datasheets; photopic response is from CIE 1931 standard; phone response is approximate for a CMOS phone sensor. (E) Smartphone user interface enables (1) patient data capture, (2) preview during focus/alignment with swipe gestures adjusting camera settings, and (3) exam data review with pinch and swipe gestures for browsing stitched image montage. (F) Photos can be uploaded directly from the smartphone to a cloud database allowing remote diagnosis with a web interface.

The device is powered by a rechargeable lithium polymer battery and includes a compact, custom-printed circuit board containing a microcontroller module (Arduino Micro, Somerville, MA), Bluetooth transceiver (RedBearLabs BLEMini, Hong Kong), two buck/boost controlled-current LED drivers, battery management controller, power supply, and several status indicator LEDs. The battery, electronics, and illumination optics are miniaturized and arranged in linear fashion along the optical axis to fit within the handle of the device (Fig. 1B). An external OLED display (4D Systems μOLED-128-G2, Minchinbury, Australia) provides a fixation target for the contralateral eye (Fig. 1B). This external display attaches to either side of the CellScope Retina housing using magnets, allowing symmetrical use while imaging either eye. Spring-loaded gold pins provide electrical contact for power and communication to the microcontroller. To conserve in-the-field battery life, the system automatically shuts off after five minutes of inactivity. Under normal daily use, the device runs for more than a week on a single charge and, like the smartphone, can be recharged with a standard USB cable.

#### Application and Software Design

A custom iPhone app was developed which communicates with the electronic hardware of CellScope Retina via Bluetooth low-energy (BLE). To begin an exam, the operator enters basic patient information and then selects an eye for imaging. The app provides a preview window which allows the operator to employ ergonomic touch and swipe motions to adjust focus, zoom, and exposure prior to initiating the image capture sequence. This allows reduction of operator motion during imaging, which can degrade image quality when using a handheld platform (Fig. 1E). Additionally, during surveying and focusing onto the retina the device uses far-red LEDs that fall outside the peak photopic response of the eye but remain within the receptive range of the iPhone camera and filters (Fig. 1C). This approach minimizes patient discomfort and unwanted gaze-induced motion artifacts. Once camera positioning and focus are optimized, the initiation of image capture signals a high-intensity white LED to flash during acquisition (Fig. 1D). The images can be uploaded to a secure server directly from the app using Wi-Fi or cellular service for remote reviewing (Fig. 1F).

The CellScope Retina hardware-software integration enables wide-field imaging of the retina in a semi-automated fashion. The device is designed to allow the operator to comfortably hold and operate the touchscreen app using one hand (Fig. 2A). The magnetically mounted screen displays a software-driven target for eye fixation to minimize unwanted eye movement during examination (Fig. 2B). This green, 2mm diameter fixation dot directs the non-imaged eye to software-specified locations on the display. Conjugate eye movements simultaneously reposition the imaged eye for rapid and precise imaging of multiple retinal fields (Fig. 2C). With the current automated program, five overlapping images are captured of the central, inferior, superior, nasal, and temporal retina in rapid sequence (Fig. 2D and Movie S4, Supplementary Information). Each image has an approximately 50-degree field-of-view and may be computationally merged with a custom algorithm that runs directly on the smartphone to create an approximately 100-degree, wide-field montage of the retina (Fig. 2E).

**Figure 2.**
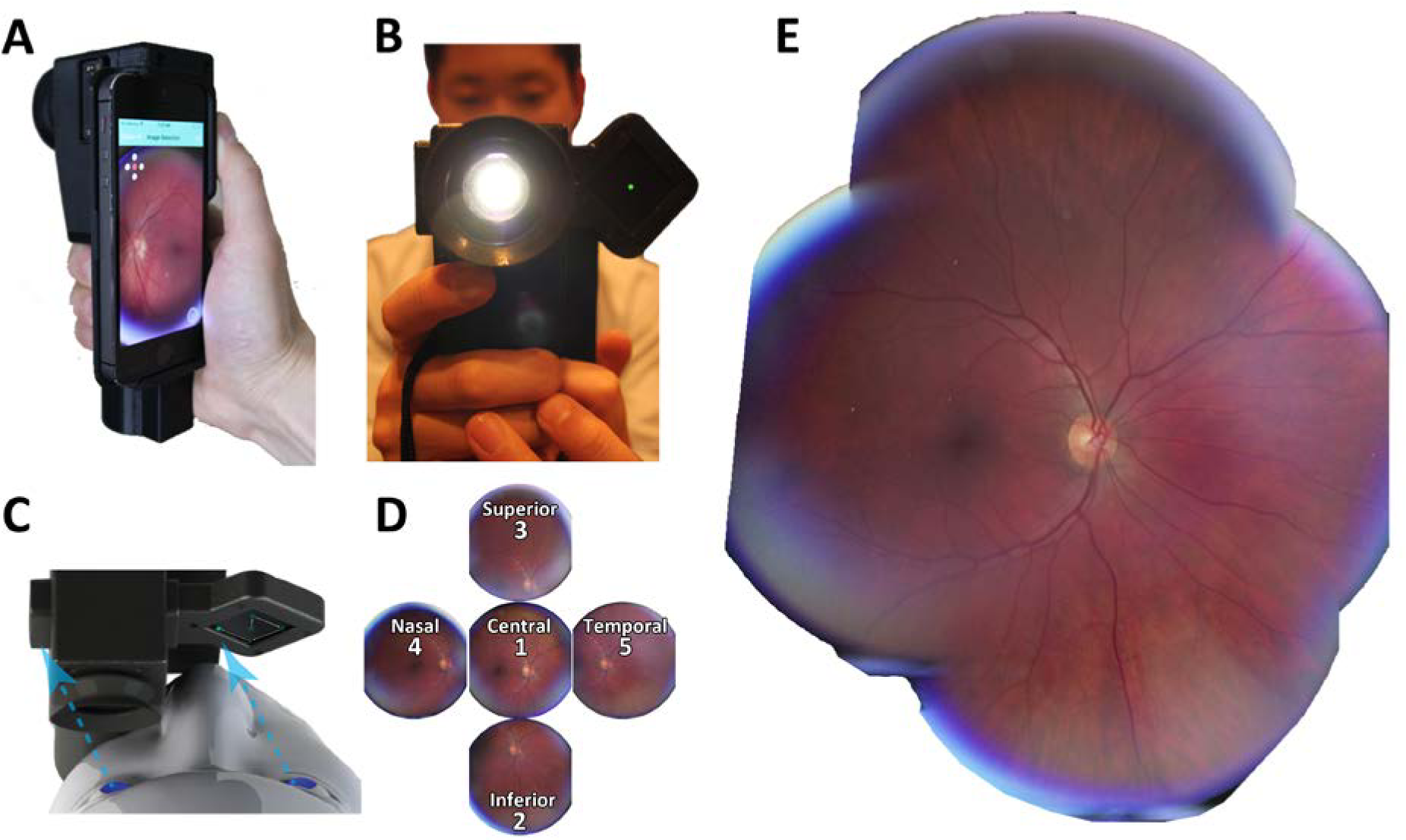
CellScope Retina is a handheld, smartphone-based retinal camera that enables fully-integrated wide-field retinal imaging. (A-B) The device is designed for easy handheld operation. (B) A detachable display may be magnetically mounted to either side of the device and provides a green dot as a fixation target. (C) The fixation target is translated through a series of positions, re-orienting the patient’s eyes and retina in a rapid and controllable fashion. (D) Each individual image is approximately 50-degrees. With the current automated arrangement, five overlapping images are captured of the central, superior, nasal, temporal, and inferior retina. (E) These five images are computationally merged on the phone using custom software to create a wide-field image of the retina spanning approximately 100-degrees.

Image-stitching could be performed either directly on the smartphone using custom software (Fig. 2E) or using the DualAlign i2k software package (Fig. 4). The custom on-phone algorithm used the OpenCV C‛‛ image-processing library for a stepwise process of correcting optical distortion, cropping the circular retinal field of view, estimating affine transforms between neighboring fields using Speeded Up Robust Features (SURF) keypoints matching. These transforms describe the translation, rotation, and skew of each peripheral field relative to the central field. To save on computational resources, the transform is estimated using down-sampled grayscale images derived from the green color channel. Finally, a full-resolution mosaic is generated using the estimated transforms and overlapping regions are linearly blended.

Processing time to generate a full resolution, 5-image montage covering approximately 100 retinal degrees (52.3 pixels/retinal degree) was ~5 minutes with our custom software on an iPhone 5s. We subsequently worked with DualAlign, LLC to generate a high-performance solution and improve on-phone processing time. A rapid preview of 100-degree montages may be produced in ~5 seconds on an iPhone 5s using downsampled images (6.4 pixels/retinal degree), allowing near real-time assessment of wide-field image data. A full-resolution montage may be generated once the operator is satisfied with image data. Initial implementation of the i2k DualAlign library has already pushed processing time for full resolution montage below 50 seconds on an iPhone 5s, with further improvements expected. All image acquisition, processing, and review steps may be performed on the device without requiring an external computer. Individual images or montages may be reviewed on the smartphone using touchscreen pinch zoom and pan gestures.

#### Study Participants

Study participants were recruited at the University of Michigan Kellogg Eye Center Retina Clinic and the ophthalmology consultation service at the University of Michigan Hospital, Ann Arbor, MI, in accordance with the University of Michigan Institutional Review Board Committee and adhered to the tenets of the Declaration of Helsinki (ClinicalTrials.gov Identifier NCT03076697). For feasibility testing in diabetic retinopathy (DR), patients (≥ 18-years-old) with diabetes and no obvious media opacity (e.g. vitreous hemorrhage, advanced cataract) were recruited while undergoing routine examination in the University of Michigan Retina Clinic. Patients in the retina clinic had a higher prevalence of retinal disease than might be expected in the general population. Images of DR were acquired with the CellScope Retina by a medical student and medical intern. For imaging of various other retinal abnormalities, patients over 18 years of age were recruited by our university-based consultation service by ophthalmology resident physicians in the emergency room or inpatient setting. A convenience sample of patients were enrolled from the consultation service based on the presence of retinal abnormalities identified from standard clinical examination. All consenting patients underwent a gold-standard dilated eye examination as part of routine care. Participants’ demographic data, clinical findings, and diagnoses were recorded.

#### Photographic and Clinical Interpretation

Patients underwent dilated fundus imaging in a dimmed room, where CellScope Retina was used to capture five sequential images (central, inferior, superior, nasal, and temporal). Both eyes were imaged unless only one eye was pharmacologically dilated for that visit. Participants were asked to assess the comfort level for red illumination and white flash from CellScope Retina (‘uncomfortable’, ‘bearable’, or ‘comfortable’), and to compare this with illumination from slit lamp biomicroscopy. CellScope Retina images were evaluated for the presence and severity of diabetic retinopathy by two masked ophthalmologists who were experienced with telemedicine grading and following review of the modified Airlie House classification grading criteria defined by the Early Treatment Diabetic Retinopathy Study (ETDRS)^33^. These graders were masked to all patient information and performed analysis per eye in a randomized fashion. Each 50-degree image was evaluated in a controlled environment on high-resolution (1600 × 1200 pixels) 19-inch displays with standard luminance and contrast on a black background in accordance with United Kingdom National Health Service guidelines^44^.

A grading template was used that assessed the quality of the image and the severity of DR. The image quality was categorized as excellent, fair, acceptable, or not gradable. DR severity was graded as mild, moderate, or severe non-proliferative diabetic retinopathy (NPDR); proliferative diabetic retinopathy (PDR); or no DR in accordance with ETDRS criteria. Because CellScope Retina acquires non-stereo images and thus cannot determine retinal thickening, photographic grading of clinically significant diabetic macular edema (CSDME) was based on the presence of hard exudates within one disc diameter of the macula center, as previously described^45, 46^. Clinically, the CSDME was defined by ETDRS criteria including retinal thickening within 500-microns of the macula center, hard exudates within 500-microns of macula center with adjacent retinal thickening, or more than one disc diameter of retinal thickening partly within one disc diameter of the macula center. Photographic diagnosis was based solely on the smartphone image acquisition, whereas clinical diagnosis was aided by Optical Coherence Tomography (OCT) and included patients actively receiving intravitreal anti-vascular endothelial growth factor therapy or intravitreal steroid therapy for CSDME.

The sensitivity and specificity in detection of referral-warranted diabetic retinopathy (RWDR) is of fundamental importance for screening programs. RWDR was defined as equal to or greater than moderate NPDR or the presence of CSDME, as previously established^33^. Traditional in-clinic diagnosis was considered the reference standard and acquired retrospectively by masked chart review of same-day visits which had included CellScope Retina imaging. In the event that the severity of DR was not explicitly documented in the chart, a masked expert reviewed the documented note, exam, and traditional imaging studies to determine disease severity. Inter-grader kappa and percent agreements were calculated to assess identical grading as well as grading within one stage. Confidence intervals for sensitivity and specificity were based on exact Clopper-Pearson confidence intervals. Statistical analysis was performed using SPSS Statistics software (IBM, Armonk, NY). Confidence intervals for the predictive values are the standard logit confidence intervals given by Mercaldo et al. 2007^47^

#### Data Availability

The datasets generated during the current study include patient information, and may be made available from the corresponding author on reasonable request complying with and approved by the relevant Institutional Review Board Committees.

### Results

#### Device

The CellScope Retina is a hand-held, battery-powered 3-D printed optical and hardware system built around a smartphone (Fig. 1). The retina is imaged via a standard ophthalmic lens and relay lenses to the cellphone camera, and the illumination and collection optical paths feature polarization-based rejection of reflections from intermediate surfaces (including the cornea), as well as far-red illumination (Fig. 1C) to increase patient comfort during focusing steps. An external side-mounted display (Figs 1A and 2B) provides a moveable fixation target to the contralateral eye to direct patient gaze (Fig. 2C); target position may be automatically stepped between positions during sequential imaging. This display is magnetically affixed, allowing switching between sides for imaging either eye. Hardware (fixation target, illumination LEDs) is controlled via Bluetooth communication with the smartphone, which runs a custom iPhone app allowing ergonomic touch and swipe motions to adjust image focus, zoom, and exposure. An automated program then collects five overlapping retinal images (Fig. 2D) which are computationally merged on the smartphone to create and display an approximately 100-degree montage of the retina (Fig. 2E). Image acquisition takes ~1 minute for cooperative patients with pharmacologic mydriasis. Multi-image montages may be generated directly on the smartphone; to our knowledge this is the first implementation of retinal montage on-phone. Our custom software produces full resolution 5-image montages with a field of view of approximately 100 degrees (52.3 pixels/retinal degree) in ~5 minutes, with image stitching running in the background on an iPhone 5s. Initial implementation of the i2k DualAlign library has already pushed this below 50 seconds, with further improvements expected. Additionally, rapid preview of 100-degree montages may be produced in ~5 seconds on an iPhone 5s using downsampled images (6.4 pixels/retinal degree). This feature allows near real-time assessment of wide-field image data during patient examination and may guide image selection for high-resolution montage. The software and mobile connectivity of the device allow images to be directly uploaded to cloud-based platforms for remote analysis or consultation.

#### Diagnostic-Quality Imaging of Various Retinal Pathologies

CellScope Retina captured diagnostic-quality images spanning a wide range of retinal findings. We demonstrate several examples including hypertensive retinopathy with a ruptured retinal arterial macroaneurysm (RAM) and hemorrhages, rhegmatogenous retinal detachment, peripapillary myelinated nerve fiber layer, circinate macular exudates, acute optic nerve edema with disc hemorrhage, non-proliferative diabetic retinopathy with dot-blot and disc hemorrhages, cytomegalovirus (CMV) retinitis, Morning Glory disc, and proliferative diabetic retinopathy with preretinal fibrovascular membrane (Fig. 3). All images were acquired after pharmacologic mydriasis of the pupil. The optics are designed such that the image nearly fills the vertical dimension of the 3:4 aspect-ratio smartphone image sensor to maximize retinal field of view in both the vertical and horizontal dimensions. Images were of sufficient quality to allow rapid diagnosis by a retinal specialist who was masked to patients’ medical records.

**Figure 3.**
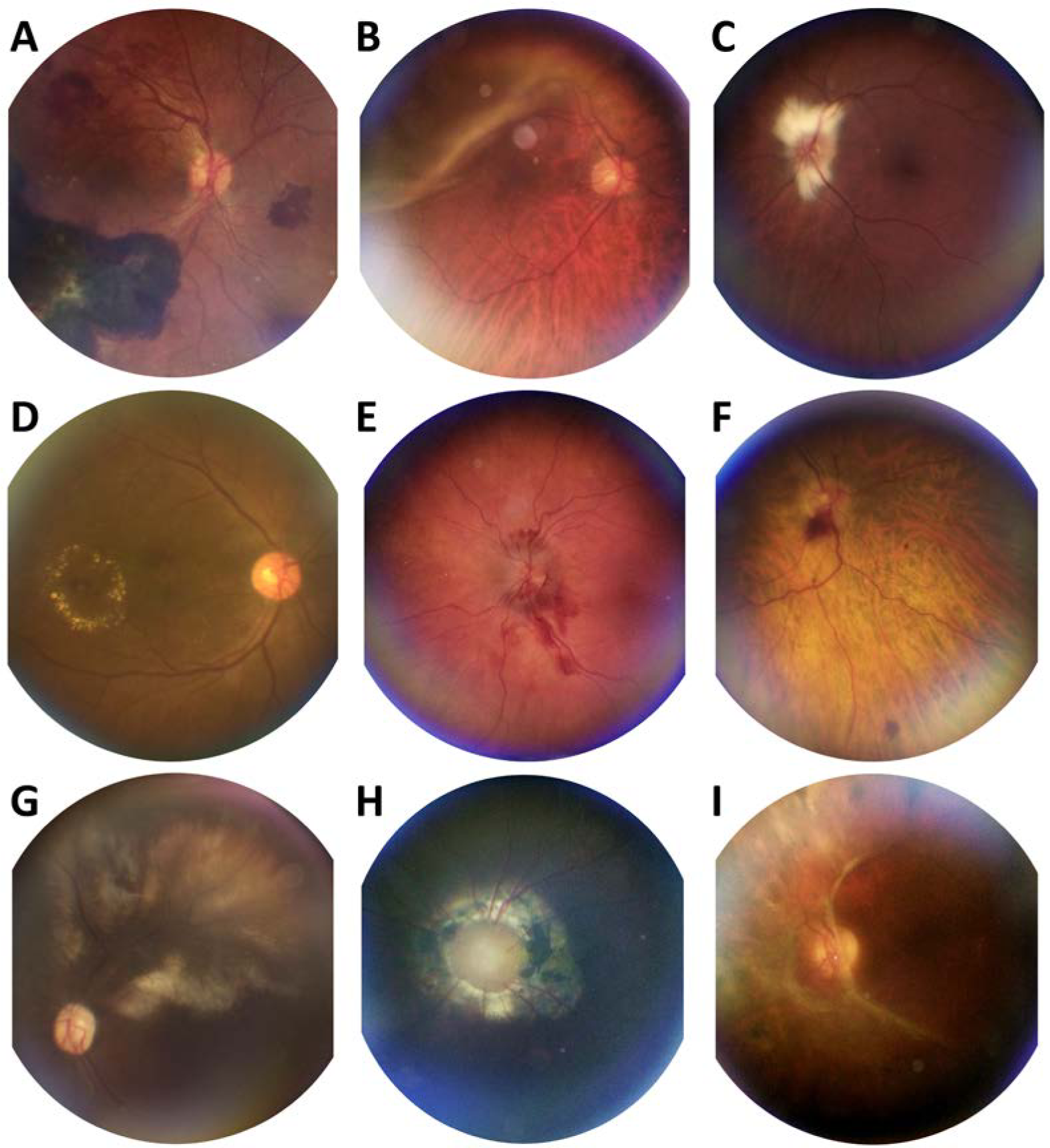
CellScope Retina produces diagnostic-quality photographs of retinal abnormalities. (A) Hypertensive retinopathy with ruptured retinal arterial macroaneurysm (RAM) along the inferior arcade with hemorrhage in multiple layers. (B) Superotemporal bullous macula-involving rhegmatogenous retinal detachment in the right eye extending from 9:00 to 1:00 o’clock. (C) Peripapillary myelinated nerve fiber layer with flame-shaped appearance and feathered borders in the left eye. (D) Circinate retinal exudates in the inferotemporal macula of the right eye. (E) Acute papilledema with flame hemorrhages extending from the optic disc and proximally along the inferior arcade of the left eye. (F) Diabetic retinopathy with dot-blot and disc hemorrhages in the right eye. (G) Cytomegalovirus (CMV) retinitis demonstrating broad retinal whitening along the superior arcade of the left eye. (H) Morning Glory disc anomaly in the left eye. (I) Epiretinal membrane with fibrovascular changes in the left eye. All images were acquired after pharmacologic mydriasis.

#### Retinal Imaging in Hospital Settings

The CellScope Retina was also evaluated outside traditional eye care facilities to assess its feasibility of use for retinal image acquisition in the inpatient and emergency department setting. Here, we demonstrate wide-field retinal photography using CellScope Retina in the inpatient setting (Fig. 4). Images of two patients with presumed fungal endophthalmitis were taken at the bedside without either repositioning the patient or using head stabilization devices. The first patient was cooperative with the examination and dilated well to 8mm with pharmacologic mydriasis, enabling 100-degree image acquisition in less than 1 minute using device-assisted fixation. Fundus photography demonstrates three creamy white, well-circumscribed 500-micron lesions within 2 mm of the optic nerve (Fig. 4A). The second patient was critically ill with altered mental status and had a poorly-dilated pupil of 4.5mm despite pharmacologic mydriasis. Multiple images were acquired over the span of approximately 4 minutes by repositioning the angle of the device relative to the eye due to the patient’s inability to fixate (Fig. 4B). Under these conditions, individual images exhibited significant glare and restricted views of the retina (Fig. 4B’). Multi-image stitching was employed to produce a >100-degree view of the retina. Fundus photography demonstrates a 16mm large, creamy-white chorioretinal lesion with associated retinal hemorrhages, retinal vasculitis, and vitritis extending from the superior arcade into the macula and threatening the fovea (Fig. 4B).

**Figure 4.**
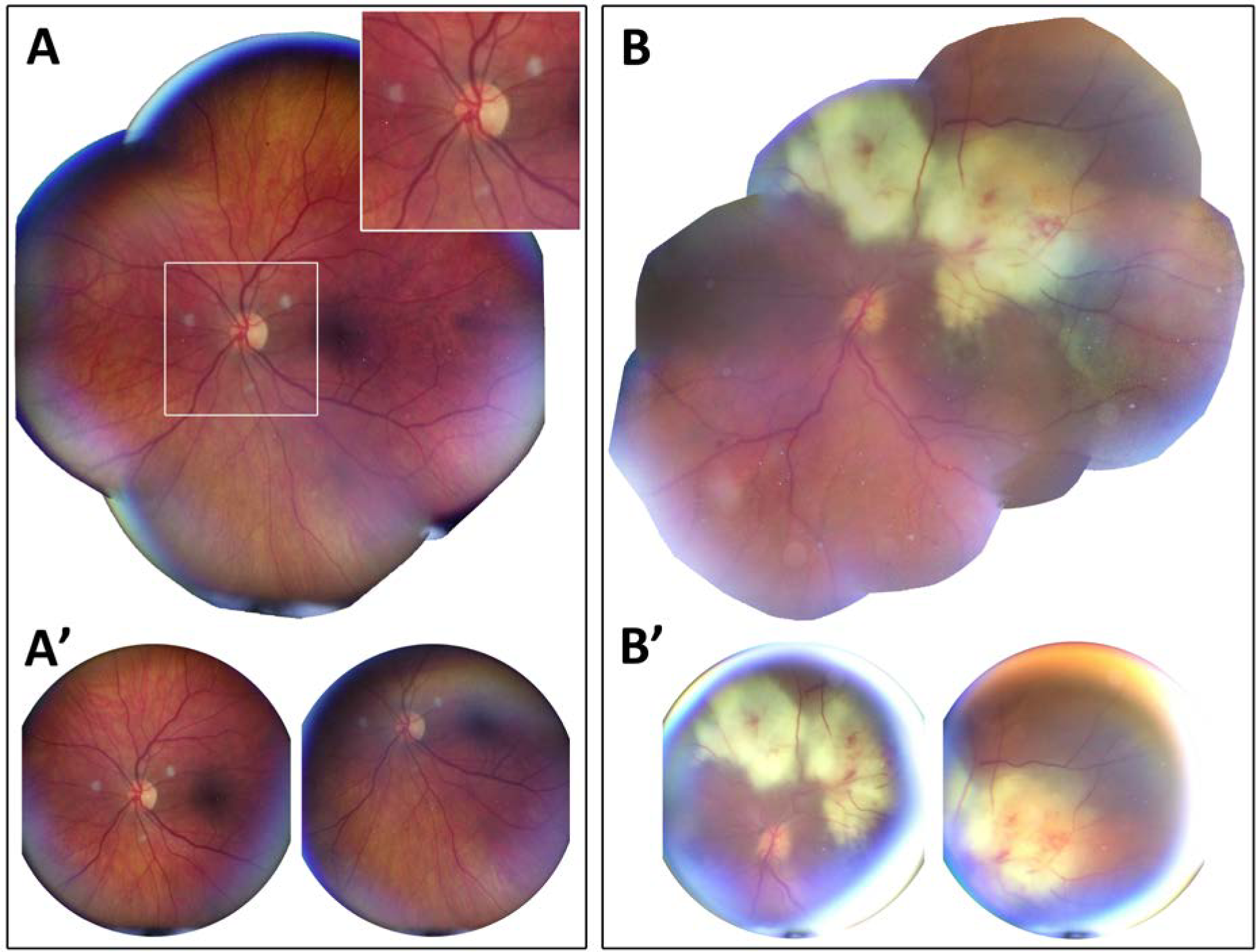
Wide-field retinal imaging with CellScope Retina under typical and challenging imaging conditions in the inpatient setting. (A) Retinal montage of presumed fungal endophthalmitis in a patient with a well-dilated 8.0mm pupil who was alert and cooperative with examination demonstrating three creamy white, well-circumscribed 500-micron lesions within 2 mm of the optic nerve. (A’) Representative images used for retinal montage. Images A and A’ were acquired in less than 1 minute by guiding the patient’s eyes with device-assisted fixation. (B) Retinal montage of presumed fungal endophthalmitis in a hospitalized patient with a poorly-dilated 4.5mm pupil and altered mental status demonstrating a 16mm large creamy white chorioretinal lesion with associated retinal hemorrhages, retinal vasculitis, and vitritis extending from the superior arcade into the macula and threatening the fovea. (B’) Representative images demonstrating glare and restricted retinal view as present with small pupil size. These images were acquired in ~4 minutes by manually repositioning the angle of the device relative to the eye due to patient’s inability to fixate. Multi-image processing steps are employed to improve wide-field viewing of the retina. All images were taken of bed-bound patients under normal inpatient care conditions without additional head stabilization or repositioning. Image stitching was performed using the i2k DualAlign software package. (Fig. 4B’).

#### Utility in Diabetic Retinopathy Screening

The CellScope Retina was further evaluated as a DR screening tool to assess its ability to resolve retinal findings in diabetic retinopathy. We demonstrate two examples of retinal imaging in patients with referral-warranted diabetic retinopathy (Fig. 5). The first patient demonstrates clear retinal abnormalities including pan-retinal photocoagulation (PRP) in both eyes. The right eye remains normal while the left eye shows re-activation of proliferative diabetic retinopathy with pre-retinal hemorrhage (Figs 5A, 5B, respectively). The second patient is diabetic without a known prior history of retinopathy. Imaging with CellScope Retina in the left eye resolves trace neovascularization and sparse microaneurysms (Figs 5D, 5E, respectively). Additionally, post-processing these images with red-subtraction was employed to enhance contrast of vascular structure within the superficial retinal layers (Figs 5D’, 5E’), an approach often used in traditional fundus photography.

**Figure 5.**
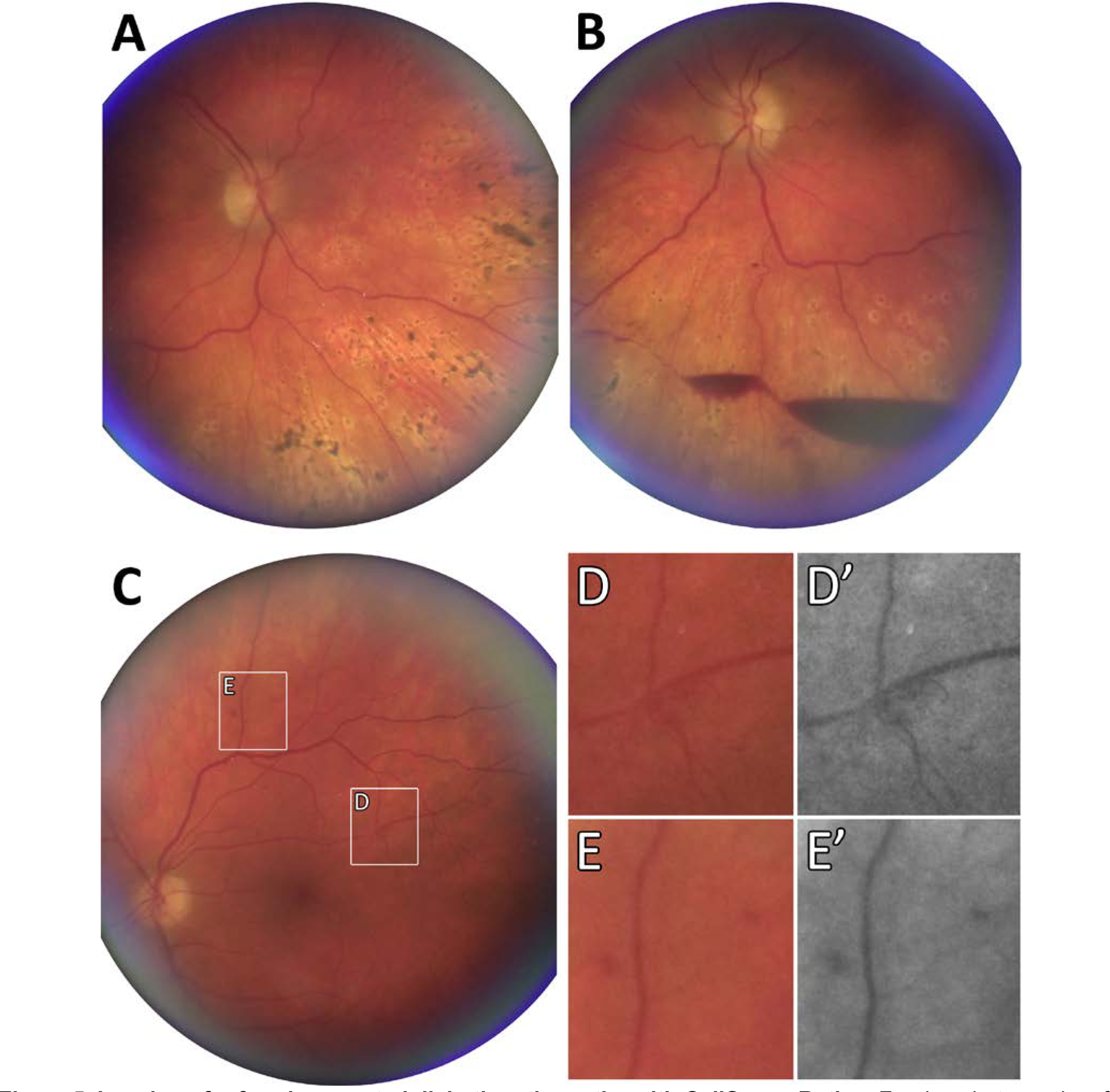
Imaging of referral-warranted diabetic retinopathy with CellScope Retina. Fundus photographs of a patient with diabetes mellitus with right (A) and left (B) eye demonstrating prior pan-retinal photocoagulation and re-activation of quiescent diabetic retinopathy with pre-retinal hemorrhage in the left eye. C) Fundus photograph of the left eye in a patient with diabetes mellitus but no known history of diabetic retinopathy. CellScope Retina resolves trace signs of neovascularization (D) and sparse microaneurysms (E). Post-processing with red-subtraction can be used to enhance contrast of vascular structures in the superficial retina, including neovascularization (D’) and microaneurysms (E’).

#### Pilot Testing for Diabetic Retinopathy Screening

In order to evaluate the potential utility of CellScope Retina for DR grading, a pilot test of 71 subjects and 142 eyes were enrolled from the University of Michigan Kellogg Eye Center Retina Clinic. 45 subjects were male, 26 were female, and the mean age was 56.7 years (standard deviation 16.9 years). CellScope Retina imaging was performed in 121 eyes (84%). The remaining 21 eyes were not imaged because patients were pharmacologically dilated in only one eye or were discharged from clinic prior to imaging both eyes. Of the 121 images evaluated by the graders, a grade of “Excellent” was given 21 times, a grade of “acceptable” was given 98 times, a grade of “fair” was given 52 times, and “ungradable” was given 2 times.

#### Detecting Referral-Warranted Diabetic Retinopathy

The sensitivity and specificity of referral-warranted DR detection by grader 1 were 91.4% (95% CI 83.0, 96.5), and 64.9% (95% CI 47.5, 79.8), respectively (Table 2). The positive and negative predictive values were 85.1% (95% CI 78.5-89.9), and 77.4% (95% CI 61.9-87.9), respectively. For grader 2, the sensitivity and specificity for referral-warranted DR detection were 95.1% (95% CI 88.0-98.7), and 48.7% (95% CI 31.9-65.6), respectively (Table 2). The positive and negative predictive values were 80.4% (95% CI 74.9-84.9), and 81.8% (95% CI 62.1-92.5), respectively.

#### Agreement between Image-Based Grading and Clinical Exam

Compared to the clinical examination, for grader 1, there was exact agreement in DR severity grading in 57.3% of the eyes, and agreement within 1 clinical stage in 88.9% of the eyes, weighted κ = 0.63 (95% CI 0.53-0.73, Table 1). For grader 2, there was exact agreement in 52.9% of the eyes, and agreement within 1 clinical stage in 81.5% of the eyes, weighted κ = 0.55 (95% CI 0.45-0.64, Table 1). The two graders agreed perfectly 56.3% of the time and were within 1 step of perfect agreement 95.0% of the time (Fig. 6), with weighted κ = 0.66 (95% CI 0.57-0.74; c.f. Table S1, Supplementary Information).

**Table 1:** Grader Assessments of Diabetic Retinopathy Severity of CellScope Retina Images versus Clinical Fundus Exam.

|  | Gold Standard Clinical Diagnosis |  |  |  |  | Weighted<br>Kappa<br>(95% CI) |
| --- | --- | --- | --- | --- | --- | --- |
|  | None | Mild<br>NPDR | Moderate<br>NPDR | Severe<br>NPDR | PDR |  |
| **Grader 1** |  |  |  |  |  |  |
| None | 11 | 5 | 0 | 0 | 3 | **0.60**<br>(0.49,<br>0.71) |
| Mild NPDR | 3 | 6 | 1 | 0 | 1 |  |
| Moderate NPDR | 1 | 6 | 7 | 1 | 2 |  |
| Severe NPDR | 0 | 5 | 10 | 6 | 10 |  |
| PDR | 0 | 1 | 0 | 1 | 37 |  |
| **Grader 2** |  |  |  |  |  |  |
| None | 1 | 3 | 0 | 0 | 0 | **0.50**<br>(0.39,<br>0.61) |
| Mild NPDR | 10 | 6 | 1 | 0 | 2 |  |
| Moderate NPDR | 3 | 9 | 12 | 3 | 9 |  |
| Severe NPDR | 1 | 4 | 3 | 3 | 3 |  |
| PDR | 0 | 1 | 2 | 2 | 41 |  |

NPDR = non-proliferative diabetic retinopathy; PDR = proliferative diabetic retinopathy.

**Table 2:** Grader Sensitivity and Specificity of Referral-Warranted Diabetic Retinopathy.

|  | Gold Standard Clinical<br>Diagnosis |  |  |  |
| --- | --- | --- | --- | --- |
| Grader Diagnosis | *RWDR* | *Non-RWDR* | *Sensitivity (95% CI)* | *Specificity (95% CI)* |
| **Grader 1** |  |  |  |  |
| RWDR | 74 | 13 | **91.4** (83.0, 96.5) | **64.9** (47.5, 79.8) |
| Non-RWDR | 7 | 24 |  |  |
| **Grader 2** |  |  |  |  |
| RWDR | 78 | 19 | **95.1** (88.0, 98.7) | **48.7** (31.9, 65.6) |
| Non-RWDR | 4 | 18 |  |  |

RWDR = referral-warranted diabetic retinopathy = equal to or greater than moderate NPDR, or the presence of CSDME

**Figure 6.**
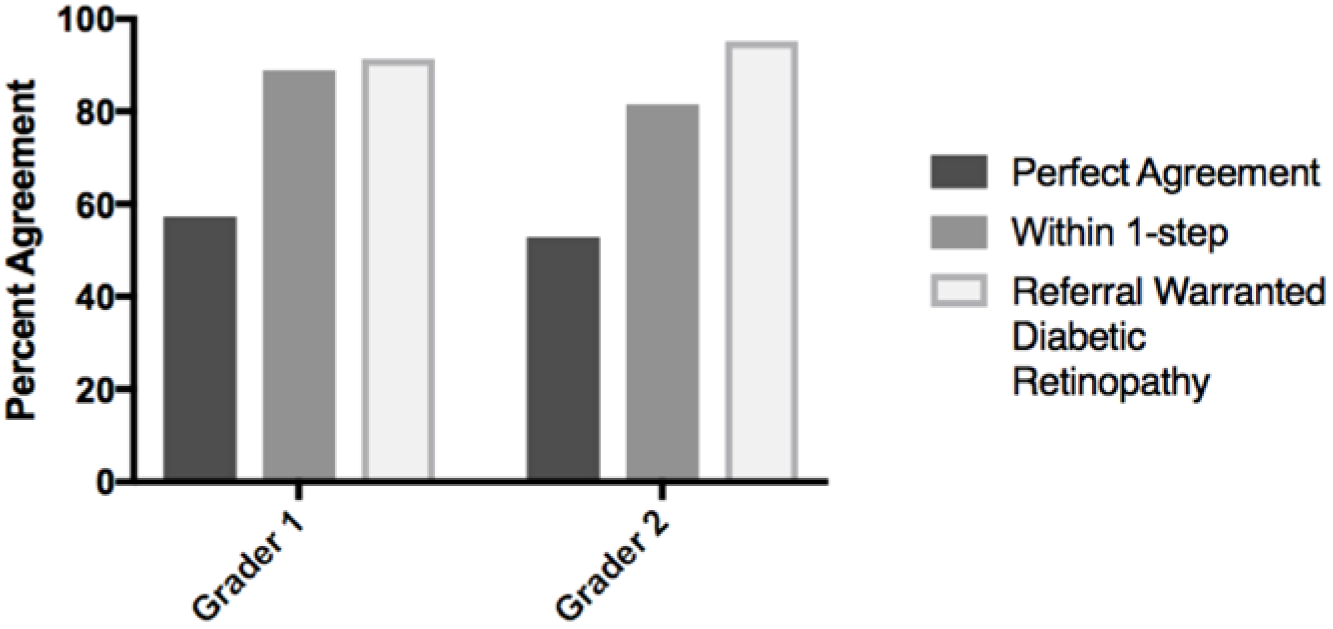
Agreement in DR severity between masked reading of CellScope Retina images and clinical examination. Diabetic retinopathy severity was determined by two masked graders and compared to grading from the clinical examination. For grader 1, there was exact agreement in 57.3% of the eyes, and agreement within 1 step in 88.9% of the eyes (kappa coefficient was 0.63, 95% CI, 0.53 - 0.73). Grader 1 was also able to detect 91.4% of referral warranted diabetic retinopathy as compared to the dilated fundus examination. For grader 2, there was exact agreement in 52.9% of the eyes, and agreement within 1 step in 81.5% of the eyes (kappa coefficient was 0. 55, 95% CI, 0.45 - 0.65). Grader 2 was also able to detect 95.1% of referral warranted diabetic retinopathy as compared to the dilated fundus examination.

#### Detecting Clinically Significant Diabetic Macular Edema

For grader 1, the sensitivity and specificity for detecting clinically significant macular edema (CSDME) was 29.4% (95% CI 10.3-56.0), and 98.0% (95% CI 92.9-99.8). The positive and negative predictive values were 71.4% (95% CI 34.5-92.2) and 89.0% (95% CI 85.6-91.7), respectively. For grader 2, the sensitivity and specificity for detecting CSDME were 68.8% (95% CI 41.3-89.0), and 83.3% (95% CI 74.4-90.2), respectively. The positive and negative predictive values were 40.7% (95% CI 28.3-54.5), and 94.1% (95% CI 88.5-97.1), respectively; c.f. Table S2, Supplementary Information. CSDME with or without DR was classified as referral-warranted disease. The in-clinical diagnosis of CSDME was assisted by use of macular optical coherence tomography.

### Patient Comfort

Patient comfort levels during examination with the red illumination used to focus on the retina (Fig. 1C) and the white flashes for image capture (Fig. 1D) were surveyed according to a three-point scale including: ‘comfortable’, ‘bearable’, or ‘uncomfortable’. All of the surveyed patients (Table S3, Supplementary Information) reported the continuous red illumination to be comfortable throughout the examination (N = 87/87). 90.8% of patients reported the white, image-capture LED flash to be comfortable (N=79/87), 8.0% reported the white flash as bearable (N=7/87), and 1.1% reported the white flash to be uncomfortable (N=1/87). Additionally, 100% of patients who also underwent slit-lamp examination reported the CellScope Retina illumination system to be more comfortable than the illumination from the slit-lamp (N=72/72).

## Discussion

Smartphone-based retinal photography holds great promise in ophthalmic care and telemedicine by leveraging the smartphone for its portability, low cost, ease-of-use, high-resolution camera, and wireless connectivity to capture diagnostic retinal images. This study highlights two important aspects of a novel, smartphone-based retinal imaging tool, the CellScope Retina. First, CellScope Retina was capable of capturing and stitching wide-field, 100-degree images of a broad variety of retinal pathologies in the non-optimal imaging conditions of an ophthalmic consultation service and emergency department setting. Second, the CellScope Retina achieved good sensitivity and moderate specificity (with high inter-grader agreement) when operated by non-ophthalmic personnel for detection of referral-warranted diabetic retinopathy in a feasibility study in this population. Furthermore, the portability and ease of one-handed operation of CellScope Retina make it possible for the device to increase access to high-quality retinal imaging when compared to benchtop fundus cameras.

There has been substantial effort in recent years to incorporate smartphone-based devices in screening and monitoring of diseases such as glaucoma and diabetic retinopathy (DR)^13–15^. In particular, three recent studies demonstrate that smartphone-based retinal imaging can be effective in screening for diabetic retinopathy^4, 16, 39^. Rajalakshmi et al. compared grading of DR from images acquired by the Fundus on Phone (FOP) system with traditional benchtop fundus photography. Using the FOP system, sensitivity and specificity for any DR was reported as 92.7% and 98.4%, respectively, and for sight-threatening DR (defined as proliferative DR or CSDME) reported as 87.9% and 94.9%, respectively^4^. Russo et al. compared grading of DR from images acquired by the D-Eye smartphone system against clinical examination with slit lamp biomicroscopy. They reported exact agreement in grading of severity of DR in 85% of cases and within 1 step of agreement in 96.7%, and a sensitivity and specificity for clinically significant macular edema as 81% and 98%, respectively^39^. Toy et al. compared grading of DR from images acquired by a smartphone with macro lens adapter and external light source compared with clinical examination. They reported that smartphone-acquired photos demonstrated 91% sensitivity and 99% specificity in detection of moderate nonproliferative DR and worse DR^16^.

While these results are encouraging, the prior literature has not addressed whether handheld retinal photography could be made simple and intuitive for non-expert operators in non-ideal settings. Indeed, experience with our first generation Ocular CellScope prototype demonstrated that smartphone imaging of the retina, while highly possible, could also be highly variable. The FOP system used an indirect ophthalmoscopy arrangement with a field-of-view of up to 45 degrees with mydriasis, which they reported could be used as a handheld system or adapted to a slit lamp frame^4^. However, the FOP system has a relatively large size and the study did not disclose who performed photography or whether the device was mounted to the slit lamp. Rigid stabilization of the camera and patient’s head helps with imaging quality but is not readily available in many community-based screening environments. The D-Eye uses a direct ophthalmoscopy arrangement with a field-of-view of up to 20-degrees with mydriasis. This requires significant manual adjustment and familiarity with retinal anatomy to perform complete retinal imaging, and images in their study were acquired by a retina specialist^39^.

Variability in image quality that could arise from non-ideal settings and non-expert operators is a critical consideration when screening in the community^37, 38^. In order to assess this potential limitation with the CellScope Retina, we performed a feasibility study wherein image acquisition for grading of DR was performed by a medical student and medicine intern rather than ophthalmologists or ophthalmic photographers. We found that even when image acquisition was performed by non-expert operators, grading of CellScope Retina images demonstrated good agreement for DR and CSDME compared with clinical examination by slit lamp biomicroscopy, consistent with other photographic-based methods^48^, and met British Diabetic Association guidelines for sensitivity in screening tools used in DR^40^. This was achieved, in part, by leveraging the processing power of the phone to simplify operational steps in image acquisition. In particular, we focused on portable, wide-field imaging due to the importance of peripheral retina surveillance in diseases such as diabetic retinopathy. Previous studies have found that a single 45-degree field of view retinal image had relatively good detection of any DR but was inadequate to effectively determine severity of DR as needed for referral^34–36^. Additionally, a prospective study using ultrawide field imaging with 200-degree field of view (Optos plc) found that approximately one-third of lesions (hemorrhages, microaneurysms, intraretinal microvascular abnormalities, neovascularization) were outside traditional ETDRS 7-field photography (90-degree field of view), and 10% of patients had a more severe grade of DR than determined by ETDRS photography^49^. These studies suggest that wider-field surveillance of the retina may improve accuracy in screening of diabetic retinopathy. At the same time, cost and speed are equally important considerations for screening in community settings. Thus CellScope Retina was designed to maximize retinal field-of-view while being fast, easy-to-use, and inexpensive.

This study has several important strengths. First, masked grading of diabetic retinopathy was performed without providing patients’ medical history, helping to eliminate bias and ensuring assessment was based purely on photographic findings. Second, non-ophthalmic operators acquired images for DR grading and, as a result, images of lesser quality were included in analysis and more closely simulated conditions that may be encountered when screening in the primary care setting. Nevertheless, CellScope Retina images both demonstrated good agreement for DR and met British Diabetic Association guidelines for sensitivity of screening tools used in DR^40^. Third, complex imaging tasks such as imaging multiple regions of the retina for wide-field analysis were simplified and standardized by leveraging computational capabilities of the phone to guide imaging. Further work in this direction has the potential to drastically simplify and improve the efficacy of retinal screening in the community.

There are several limitations to this study. First, participants for the DR study were recruited from the retina clinic in a tertiary care eye hospital, where the prevalence of patient DR is much higher than in the general population. While our feasibility study shows promising results, additional work is required to validate the accuracy and utility of the CellScope Retina in the primary care and community-based settings where the technology is most needed. Second, CellScope Retina is currently designed as a mydriatic device that requires patients’ eyes to be pharmacologically dilated. This can be time-consuming and uncomfortable for patients while also being unfamiliar to primary care physicians. Third, a medical student and a medicine intern acquired the images for the DR study, and additional validation for ease of imaging would be called for if shifting to use by medical assistants and primary care personnel.

Smartphone-based retinal imaging offers the possibility of dramatically increasing the accessibility of ophthalmic care by remote screening and diagnosis of vision threatening diseases among millions of individuals not receiving regular eye examinations. Achieving this goal will rely not only on the familiarity and portability of smartphones, but also on the development of novel techniques that enhance image quality and reliability. We aimed to make wide-field retinal imaging intuitive and reproducible for the non-expert operator. The resulting CellScope Retina device harnesses hardware and software automation controlled by a smartphone to simplify retinal imaging, potentially improving access to ophthalmic care through handheld and portable retinal disease screening and diagnosis.

## Additional information

Supplementary information follows this paper, and a supplementary video is also available online. Correspondence and requests for materials should be addressed to D.A.F.

## Supplementary Information

**Supplementary Table S1:**
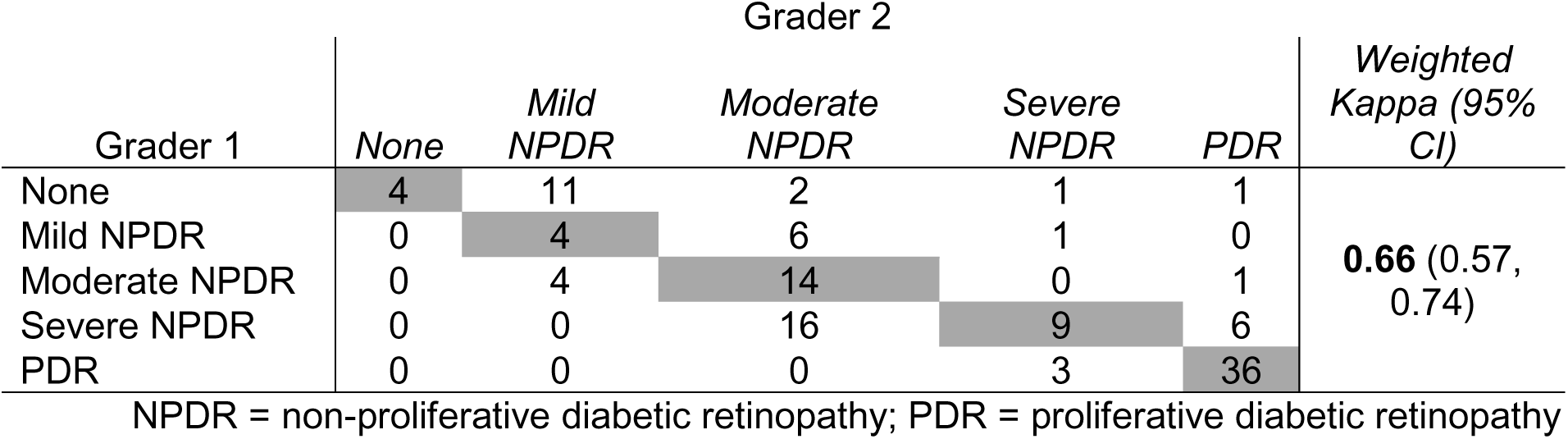
Assessment of Diabetic Retinopathy Severity Agreement between Graders.

| Grader 1 | Grader 2 |  |  |  |  | Weighted Kappa (95% CI) |
| --- | --- | --- | --- | --- | --- | --- |
|  | None | Mild NPDR | Moderate NPDR | Severe NPDR | PDR |  |
| None | 4 | 11 | 2 | 1 | 1 | **0.66** (0.57, 0.74) |
| Mild NPDR | 0 | 4 | 6 | 1 | 0 |  |
| Moderate NPDR | 0 | 4 | 14 | 0 | 1 |  |
| Severe NPDR | 0 | 0 | 16 | 9 | 6 |  |
| PDR | 0 | 0 | 0 | 3 | 36 |  |

NPDR = non-proliferative diabetic retinopathy; PDR = proliferative diabetic retinopathy

**Supplementary Table S2:** Grader Assessments of Macular Edema of CellScope Retina Images versus Clinical Fundus Exam.

|  | Gold Standard Clinical<br>Diagnosis |  |  |  |
| --- | --- | --- | --- | --- |
| Grader Diagnosis | CSDME | No CSDME | Sensitivity (95% CI) | Specificity (95% CI) |
| **Grader 1** |  |  |  |  |
| CSDME | 5 | 2 | **29.4** (10.3, 56.0) | **98.0** (92.3, 99.8) |
| No CSDME | 12 | 97 |  |  |
| **Grader 2** |  |  |  |  |
| CSDME | 11 | 16 | **68.8** (41.3, 89.0) | **83.3** (74.4, 90.2) |
| No CSDME | 5 | 80 |  |  |

CSDME = Clinically significant edema

**Supplementary Figure S3:**
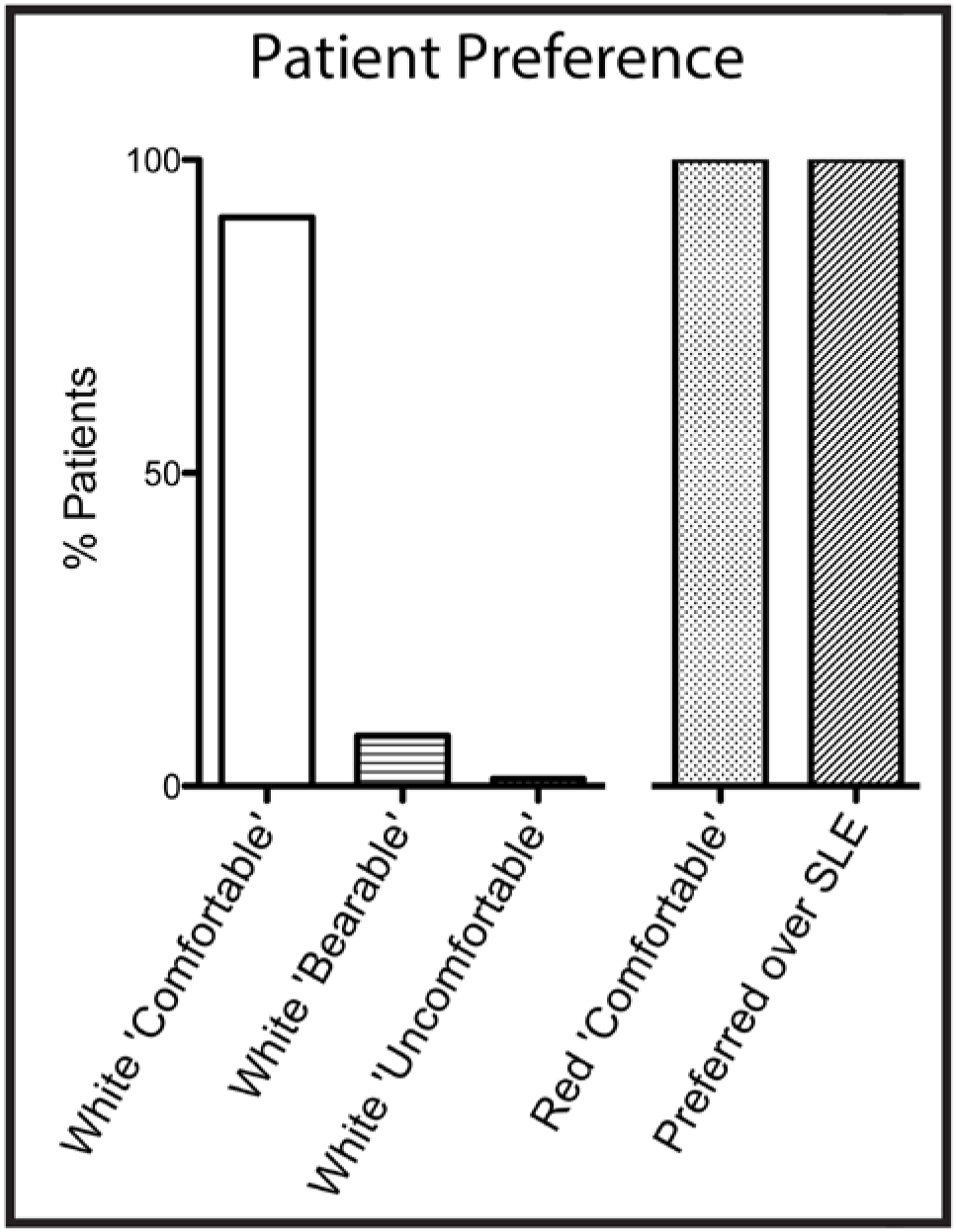
Continuous red illumination with white flash is comfortable for patients. Red illumination is used for focusing and alignment on the retina with white flash for image capture. Of surveyed patients, 90.8% reported white flash with CellScope Retina was ‘comfortable’ throughout examination (N=79/87), 8.0% reported white flash was ‘bearable’ (N=7/87), and 1.1% reported white flash was ‘uncomfortable’ (N=1/87). 100% of patients reported continuous red illumination with CellScope Retina was ‘comfortable’ for the duration of examination (N=87/87). 100% of surveyed patients that also underwent slit-lamp examination reported that illumination from CellScope Retina was more comfortable than illumination from the slit-lamp (N=72/72).

**Supplementary Movie S4, Caption:**

**CellScope Retina imaging sequence viewed from the patient perspective**. The red illumination annulus (center) is focused onto the peripheral cornea. With a properly positioned eye, the annulus is defocused to evenly illuminate the retina. The magnetically-attached fixation display can be switched to either side of the instrument. An image-capture sequence cycles a 2mm diameter green fixation dot through different positions, orienting the patient’s gaze to allow sequential imaging of the central, inferior, superior, nasal, and temporal retinal fields. Illumination for focusing is deep red (655nm) and images are acquired using white flash. Five 50-degree fields are captured and then computationally merged to create an approximately 100-degree montage of the retina.

